# Simultaneously inferring T cell fate and clonality from single cell transcriptomes

**DOI:** 10.1101/025676

**Authors:** Michael J.T. Stubbington, Tapio Lönnberg, Valentina Proserpio, Simon Clare, Anneliese O. Speak, Gordon Dougan, Sarah A. Teichmann

**Affiliations:** European Molecular Biology Laboratory, European Bioinformatics Institute (EMBLEBI), Wellcome Genome Campus, Hinxton, Cambridge, CB10 1SD, UK; Wellcome Trust Sanger Institute, Wellcome Genome Campus, Hinxton, Cambridge, CB10 1SA, UK

## Abstract

The heterodimeric T cell receptor (TCR) comprises two protein chains that pair to determine the antigen specificity of each T lymphocyte. The enormous sequence diversity within TCR repertoires allows specific TCR sequences to be used as lineage markers for T cells that derive from a common progenitor. We have developed a computational method, called TraCeR, to reconstruct full-length, paired TCR sequences from T lymphocyte single-cell RNA-seq by combining existing assembly and alignment programs with a “synthetic genome” library comprising all possible TCR sequences. We validate this method with PCR to quantify its accuracy and sensitivity, and compare to other TCR sequencing methods. Our inferred TCR sequences reveal clonal relationships between T cells, which we put into the context of each cell’s functional state from the complete transcriptional landscape quantified from the remaining RNA-seq data. This provides a powerful tool to link T cell specificity with functional response in a variety of normal and pathological conditions. We demonstrate this by determining the distribution of members of expanded T cell clonotypes in response to *Salmonella* infection in the mouse. We show that members of the same clonotype span early activated CD4+ T cells, as well as mature effector and memory cells.

## INTRODUCTION

T lymphocytes are crucial to the adaptive immune system. Key to their function is the ability of each T cell to recognise specific peptide–major histocompatibility complex (pMHC) combinations presented on the surface of antigen presenting cells^1^. This specific recognition is mediated by the T cell receptor (TCR), an extremely diverse cell-surface protein that has been estimated to have up to 10^15^ possible variations^2^. The TCR is a heterodimer of an α and a β chain that are encoded by genes produced by V(D)J recombination of two separate germline loci during T cell development^3^. The enormous nucleotide sequence diversity of paired TCR sequences allows us to assume that cells with identical paired TCR genes arose from the same T cell clone.

The diversity of single TCR chains (typically β) has been used as a proxy for overall clonal diversity within bulk populations of T lymphocytes^4–6^ but studies of bulk populations cannot determine the paired heterodimeric chains within each cell. This limits their ability to perform high-resolution determination of the clonal relationships between individual cells and also to draw conclusions about the antigenic specificities of the T cells in the studied population^1^.

An ability to study the paired TCR sequences within individual T cells will be extremely powerful in understanding the adaptive immune response in a variety of normal and pathological conditions. Knowledge of the paired TCR chains that are involved in T cell responses to particular pathogens and antigens will be crucial in informing studies intended to discern the ‘grammar’ of TCR recognition or to design therapeutic TCR molecules. Furthermore, making a connection between TCR sequence and the transcriptional identity of individual T cells will be even more informative, enabling us to connect cellular transcriptional fate with antigen specificity, to measure and model the dynamics of clonal expansion within T cell populations and to investigate T cell phenotypic plasticity.

Paired TCR analysis has been performed in individual single-cells but these methods have, until now, relied upon specific amplification of the TCR loci^7–10^ or capture of the TCR genes^11^ and so provide no other information about the cells in question. In addition, biases in PCR primer efficiency prevent accurate determination of TCR expression levels. A method that also amplifies a small set of ‘phenotyping marker’ genes^12^ provides limited information about the functional identity of the cells and requires *a priori* knowledge of genes whose expression will be informative, as well as the design and optimisation of many multiplexed PCR primers.

Single-cell RNA-seq (scRNA-seq) has already proved valuable in investigating the transcriptional heterogeneity and differentiation processes of several cell populations^13–19^ and has provided insights into a novel T lymphocyte subset^20^. However, it has not yet been possible to determine recombined TCR sequences from T cell scRNA-seq datasets.

Existing computational tools for the analysis of TCR sequences are designed for use with experiments that analyse bulk cell populations and also require the targeted amplification of TCR loci during the experimental step^21–25^. Therefore, the nature of bulk TCR-sequencing data is quite different from single-cell whole transcriptome RNA-sequencing data, and the requirements for alignment methods are distinct. In future, combining these methods with single cell data could be a fruitful strategy to gain a global overview of clonal amplifications of individual loci together with in-depth phenotyping of several hundreds to thousands of individual cells.

Here, we present a novel method and software tool that enables full-length, paired TCR sequences (alongside their expression levels) to be reconstructed from single-cell RNA-seq data with high accuracy and sensitivity. Importantly, this method requires no alterations to standard scRNA-seq protocols and so can be easily applied to any species and sample for which scRNA-seq is possible. This novel approach links clonal ancestry and antigen specificity with the comprehensive transcriptomic identity of each studied T cell.

## RESULTS

### Extraction and assembly of complete TCR sequences from RNA-seq data

We have extended the analysis of RNA-seq data from single T lymphocytes to enable the accurate and specific reconstruction of full-length sequences of recombined and expressed TCR loci within each cell. Importantly, this approach does not require any alterations to standard single-cell RNA-seq protocols and, furthermore, provides accurate, unbiased measurements of expression for thousands of genes within the transcriptome of each cell as well for the TCR loci themselves. Here, we use scRNA-seq data generated using the SMART-Seq protocol^26^ performed using the Fluidigm C1 microfluidics system. Our method would, however, work with any RNA-seq data derived from full-length cDNA.

Our method begins by extracting TCR-derived sequencing reads from the pool of all RNA-seq reads for each single cell (**Fig. 1a**). This is achieved by aligning the sequencing reads against ‘synthetic genomes’ comprising all possible combinations of V and J segments (**Fig. 1b**). A separate synthetic genome and alignment step was used for the alpha and beta TCR loci. The alignment step is performed with low stringency to ensure that the maximum number of TCR-derived reads are captured. We then assemble the potential TCR-derived reads for each locus into contiguous sequences (contigs) using the *de novo* RNA-seq assembly package, Trinity^27^. This assembly step provides resolution of sequencing errors to provide highly-accurate contig sequences.

**Figure 1.**
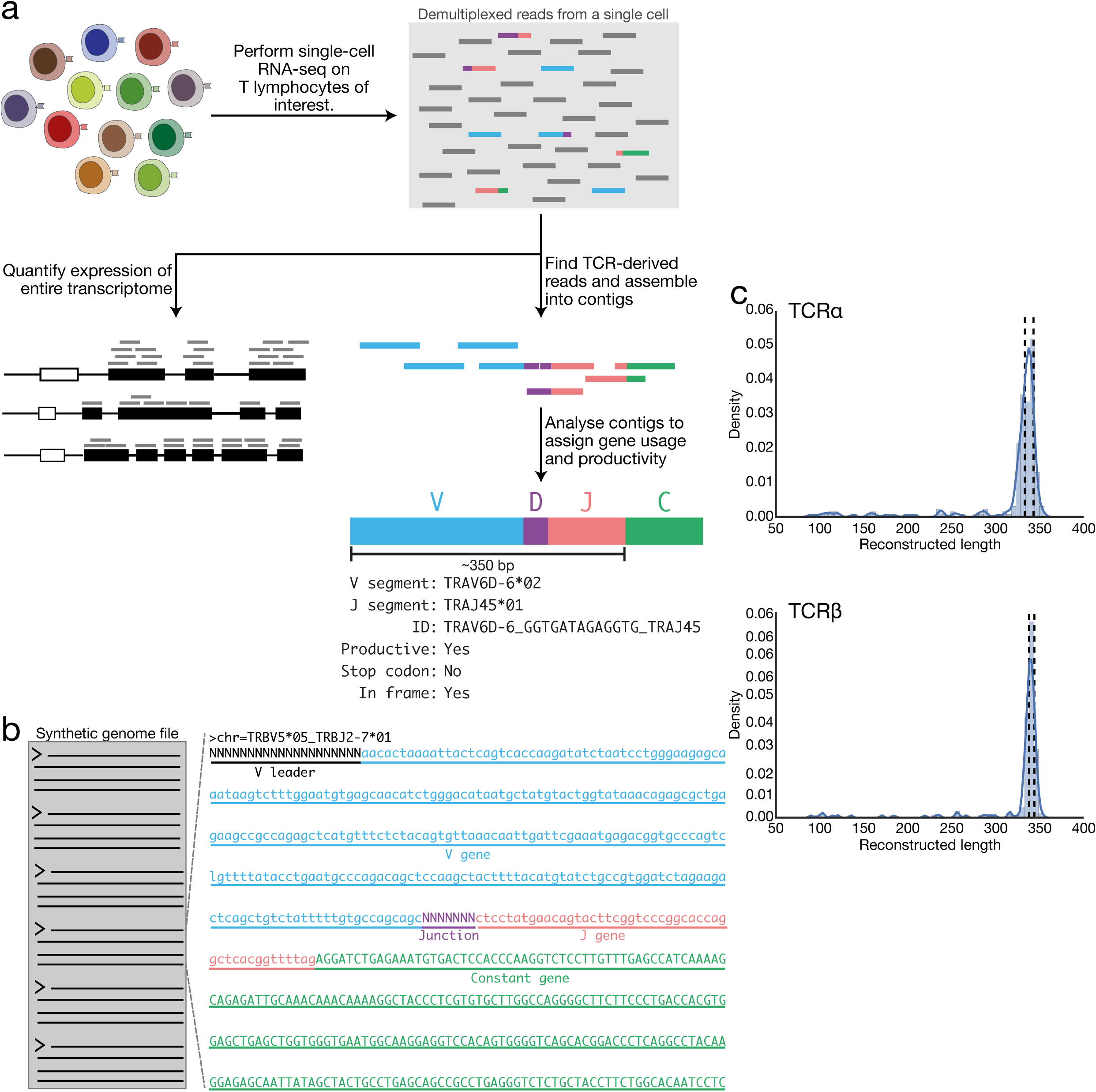
Method for reconstructing TCR sequences from single-cell RNA-seq data. (**a**) Overview of data-processing steps for TCR sequence reconstruction. Single-cell RNA-sequencing was performed on individual T lymphocytes to produce a pool of paired-end sequencing reads for each cell. These reads were used to quantify gene expression within each cell. In addition, sequencing reads that are derived from TCR mRNA are extracted and assembled into long contiguous TCR sequences. TCR contigs are filtered and analysed with IgBlast to determine the gene segments used and the junctional nucleotides. (**b**) Example of a synthetic genome entry used as alignment reference for extraction of TCR-derived reads. Each TCR locus is represented by a fasta file containing entries comprising every possible combination of V and J genes for that locus. V–J combinations contain the sequence of the appropriate constant gene along with stretches of N nucleotides to represent V leader and variable junctional regions. (**c**) Distributions of lengths of reconstructed TCR sequences. Reconstructed sequences were trimmed to include the region derived from the V gene, junction and J gene. The lengths of these sequences are plotted as histograms and kernel density estimates. Dotted lines represent the interquartile range of lengths of full-length sequences derived from the synthetic genome files.

To determine which of the assembled contigs represent full-length, recombined TCR sequences we use IgBLAST^28^ to analyse each contig and, where possible determine the V, D and J segments within the contig and the exact nucleotide sequences of the junctions. Contigs are taken forwards for further analysis if they have gene segments from the expected locus (i.e. TCRα genes for TCRα contigs) along with low IgBlast E-values indicative of high-quality alignment.

Importantly, the reconstructed recombinant sequences typically contain nearly the complete length of the TCR V(D)J region (**Fig. 1c**) and so allow high-confidence discrimination between closely related and highly-similar gene segments. Where multiple contigs from a single cell all represent the same original recombinant TCR sequence, they are collapsed into a single assignment. TCR sequences are then analysed for the presence of an appropriate open reading frame to determine whether they can produce a full-length polypeptide chain. Finally, the full nucleotide sequence of each recombinant is reduced to a recombinant identifier (**Fig. 1a**) that uses IMGT gene nomenclature^29^,^30^ and uniquely represents the V and J segments as well as the junctional nucleotides present in the sequence.

In very few cases (5/272, 1.8% of cells in this study), a cell will have more than two contigs for one or both of its TCR loci. These biologically implausible situations are resolved by quantifying the expression of each possible contig within the cell and taking the two most highly-expressed recombinants. TCR expression is quantified by appending the contig sequences to the entire mouse transcriptome and then using ‘pseudoalignment’-based abundance quantification performed by the Kallisto algorithm^31^.

To permit broad use of our approach by other researchers, we have made the TCR reconstruction tool, ‘TraCeR’, available at www.github.com/teichlab/tracer. This performs TCR sequence reconstruction as well as summarising and visualising data from multiple single cells.

### Performance, validation and comparison with PCR-based TCR sequencing

To demonstrate and validate our method for TCR sequence reconstruction we analysed single-cell RNA-seq data from 272 FACS-sorted CD4^+^ T cells isolated from spleens of C57BL/6 mice (**Table 1, Supplementary Fig. 1**). We were able to detect at least one productive alpha chain in 74%–96% of cells, a productive beta in 88%–96% and paired productive alpha-beta chains in 70%–93% (**Table 2**, **Supplementary Table 1**). This compares favourably with previous PCR-based approaches for single-cell TCR sequencing that were able to detect productive, paired TCR genes in 50%–82% of cells^7–9^,^12^.

**Table 1.**
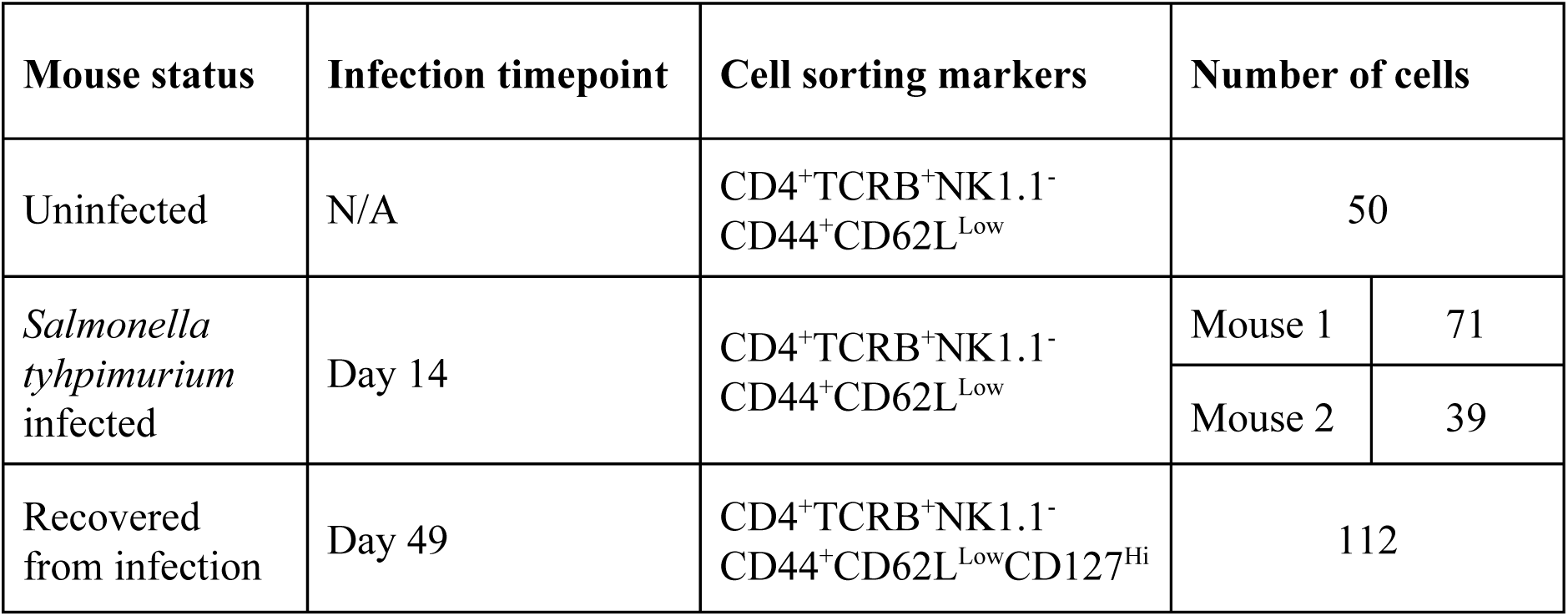
Mouse splenic CD4^+^ T lymphocyte populations used for single-cell RNA-seq

**Table 2.**
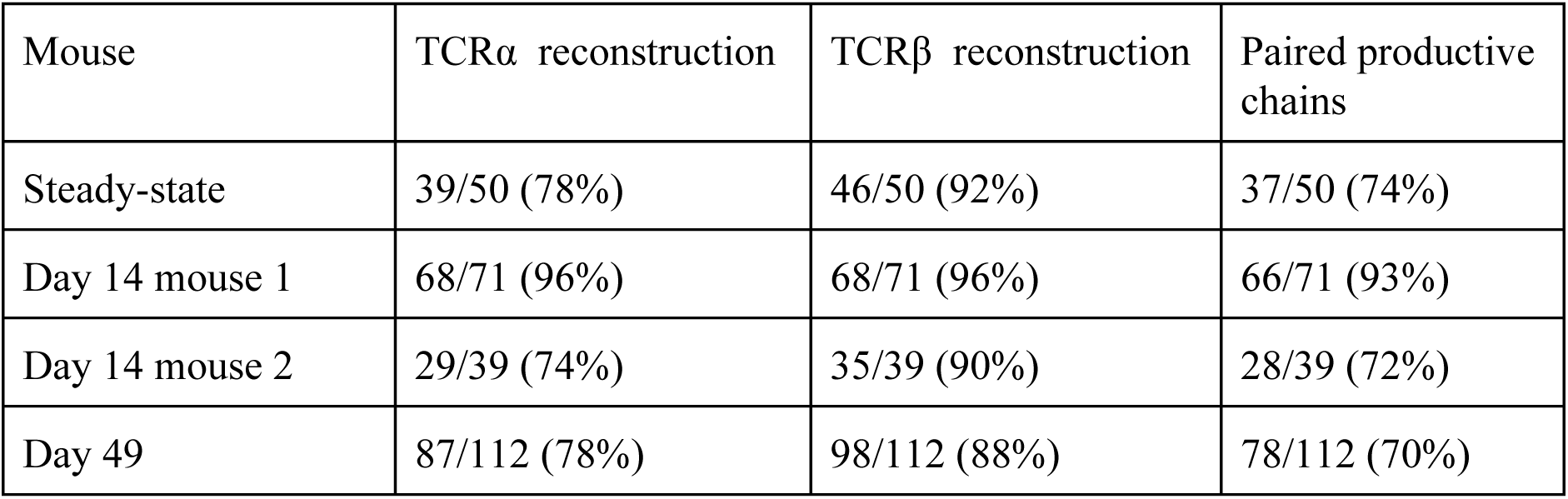
TCR reconstruction statistics

| Mouse | TCRα reconstruction | TCRβ reconstruction | Paired productive chains |
| --- | --- | --- | --- |
| Steady-state | 39/50 (78%) | 46/50 (92%) | 37/50 (74%) |
| Day 14 mouse 1 | 68/71 (96%) | 68/71 (96%) | 66/71 (93%) |
| Day 14 mouse 2 | 29/39 (74%) | 35/39 (90%) | 28/39 (72%) |
| Day 49 | 87/112 (78%) | 98/112 (88%) | 78/112 (70%) |

Many T cells contain two recombined alleles for one or both TCR chains. In the majority of cases, one allele encodes a full-length chain whilst the other is non-productively rearranged and contains a frame-shift and/or stop codons. However, it has also been shown that 30% of murine T lymphocytes possess two productively rearranged TCRα genes whilst up to 10% possess two productive TCRβ genes^32^.

Our method detected two alpha chain recombinants in 42% of cells and two beta chain recombinants in 22% (**Supplementary Table 1**). We detect two productive alpha chains in 35 out of 188 (19%) cells with at least one productive alpha chain and two productive beta chains in 16 out of 247 (6%). These data are in line with the previously reported statistics above, and suggest that our method is more sensitive than the best-performing PCR-based method, which surprisingly, did not detect multiple β recombinants in any of the 1268 cells studied^12^. In only one cell (0.3%) did we detect two non-productive sequences for a locus and both of these sequences were validated by the PCR-based approach described below.

We compared the TCR sequences reconstructed by our method to those detected by a multiplex PCR-based approach^12^ that we adapted for use with mouse cells. We took 185 cells for which we had RNA-seq data, and used full-length SMART-seq-derived cDNA as template material in PCR reactions. We designed the PCR to amplify a region from within the V gene up to the constant segment. The products from each cell also contained a unique indexed barcode (**Fig. 2a**). PCR products were purified, pooled and sequenced using an Illumina MiSeq. Sequencing reads from individual cells were then resolved according to their barcode sequences and read-pairs were merged based on their overlapping sequence.

**Figure 2.**
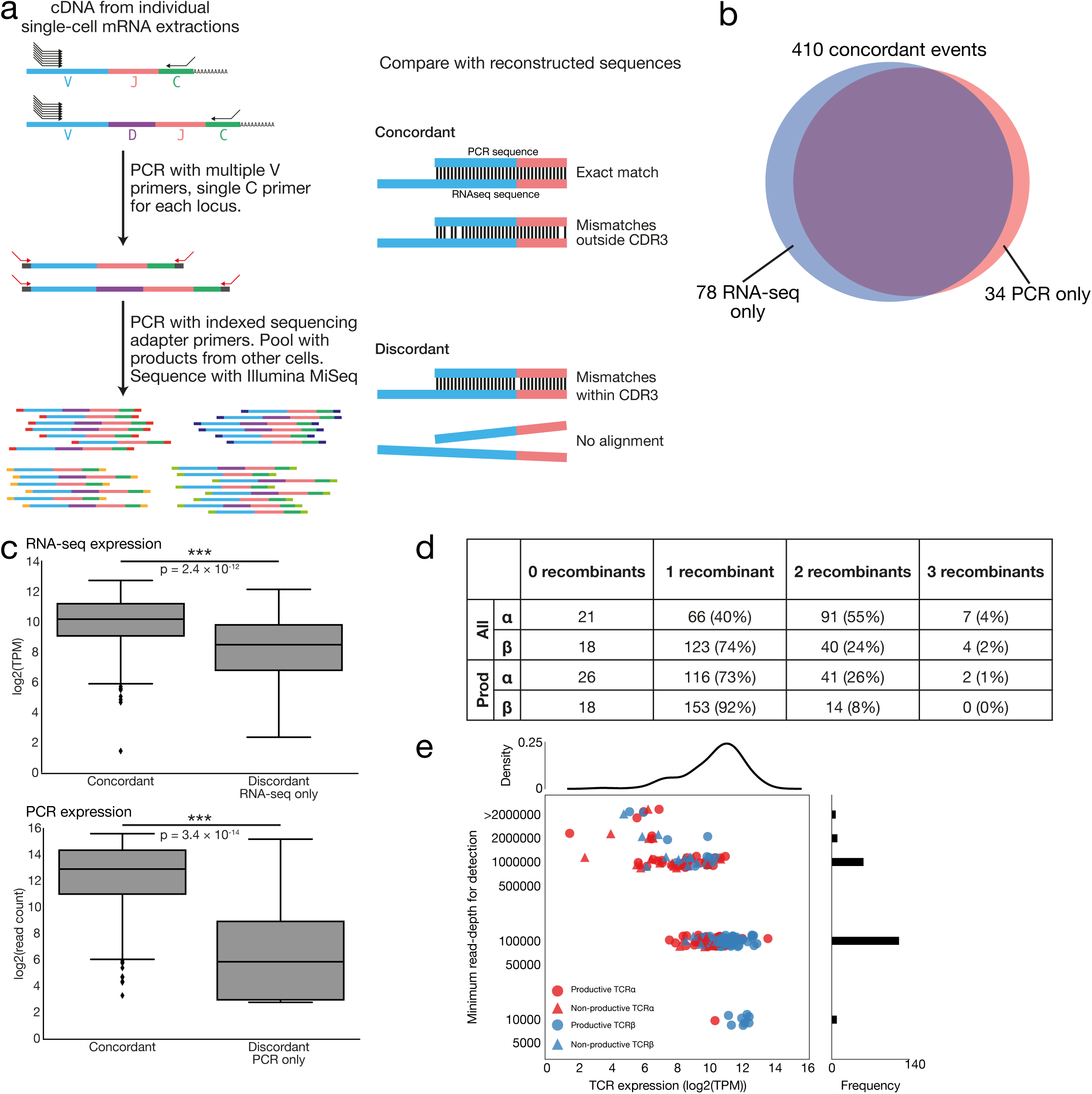
Validation of RNA-seq TCR reconstruction. (**a**) PCR amplification and sequencing of recombined TCR genes from single-cells adapted from a previously reported method^12^. Full-length single-cell cDNA libraries were used in PCR reactions with multiplexed V gene primers and single constant region primers for each TCR locus. Amplicons were purified, pooled and sequenced using an Illumina MiSeq. Sequences were compared with TCR sequences reconstructed from RNA-seq data for the same cells. Alignments were classified as concordant if there were no mismatches or if mismatches were found only outside the CDR3-encoding region. (**b**) Numbers of concordant and discordant events from comparison between RNA-seq and PCR. Concordant events include 39 occasions where no sequence was detected by either method for a particular locus. (**c**) Expression levels of concordant and discordant recombinant sequences. Expression levels of TCR sequences were calculated as transcripts per million (TPM) from RNA-seq data or as numbers of reads from PCR data. P values were calculated using the Mann-Whitney U test. (**d**) Number of cells with zero, one, two or three recombinants for each TCR locus from combined RNA-seq and PCR results. Either for all (‘All’) or productive recombinants only (‘Prod’). (**e**) Sensitivity analysis of RNA-seq reconstruction. All single-cell datasets from day 14 mouse 1 were randomly subsampled three independent times to contain decreasing total read numbers followed by TCR reconstruction. Points representing each TCR sequence found in the full datasets are plotted according to their expression levels and the minimum total read depth required for detection in at least two out of three subsamples. For clarity, points are jittered about the y-axis.

Sequences with up to 5% similarity were merged and a consensus was derived to overcome sequencing and PCR errors. Sequences were then analysed by IgBlast to determine if they were from rearranged TCR genes and were filtered to remove background sequences that were not supported by sufficient sequencing reads.

For each cell, TCR sequences derived from PCR amplification were aligned against those reconstructed from the RNA-seq data to determine whether a particular recombinant had been detected by both methods. The number of mismatches throughout the entire alignment and also within the CDR3-encoding region were determined (**Fig. 2a**). Alignments that differed within the CDR3 region were considered to be derived from different original sequences. Mismatches within the remainder of the alignment were permitted since these were likely to be due to errors introduced during PCR amplification or sequencing.

The alignments between RNA-seq and PCR-derived TCR sequences were used to classify each recombinant in each cell as concordant (detected by both approaches) or discordant (detected by only one approach). In total, 485 recombinant sequences were detected by one or both approaches, of which 371 (76.5%) were concordant (**Supplementary Table 2**). From the 371 concordant sequences, 55 had at least one mismatch throughout the non-CDR3 portion of the alignment with, at most, three mismatches occurring.

In addition, there were 39 occasions where no sequence was detected by either technique for a particular locus, thereby giving 410/523 (78.4%) concordant events (**Fig. 2b**). Thirty-five recombinant sequences (6.7%) were detected only by PCR whilst 79 (15.1%) were present only in the RNA-seq data.

To determine the cause of the discordant recombinants, we first investigated whether they were, in fact, artefacts caused by misassembly or sequencing errors. We checked whether discordant recombinant sequences within a particular cell were highly similar to other sequences (discordant or concordant) that were also detected within the same cell. We only found one example of this (**Supplementary Fig. 2**) and, here, the difference was within a long homopolymeric stretch (11 or 13 nt) of G nucleotides. Long homopolymeric regions are problematic for PCR amplification and sequencing reactions so it is difficult to draw conclusions as to which sequence represents the actual TCR within the cell.

We then hypothesised that discordant reads might represent genuine TCR sequences present within the cells that are expressed at lower levels. These will be less likely to be concordantly detected by both approaches due to stochastic detection of sequences expressed at levels approaching the sensitivity limits of each technique. We found that discordant recombinants are, indeed, expressed at lower levels than concordant recombinants (**Fig. 2c**). Further evidence for this hypothesis is provided by determination of the numbers of total recombinants and productive recombinants within each cell if the RNA-seq and PCR data are combined (**Fig. 2d**). Only 10/185 (5%) cells contain more than two recombinants for a locus, which is biologically implausible, whilst the numbers of cells with two productive recombinants for each locus are in accordance with previous reports^32^.

Finally, we sought to determine the number of sequencing reads required for each cell to permit detection of recombinant sequences. To do this we randomly subsampled sequencing reads from all the cells from a single mouse (day 14 mouse 1). Reads were subsampled three independent times to depths of 2, 1, 0.5, 0.1, 0.05, 0.01 and 0.005 million paired-end reads. We then used TraCeR to attempt reconstruction of the TCR sequences and determined in how many subsamples at each depth the expected recombinants were detected. We found that more highly-expressed TCR genes could be reconstructed at lower read depths and that 92% of recombinants were reliably reconstructed from 1 million reads or fewer (**Fig. 2e, Supplementary Fig. 3**). Therefore, our method does not require sequencing depth above that which is optimal for accurate transcript quantitation (Power Analysis of Single Cell RNA-Sequencing Experiments. Svensson V., Labalette C., Macauley IM., Cvejic A., SAT. *Submitted*).

**Figure 3.**
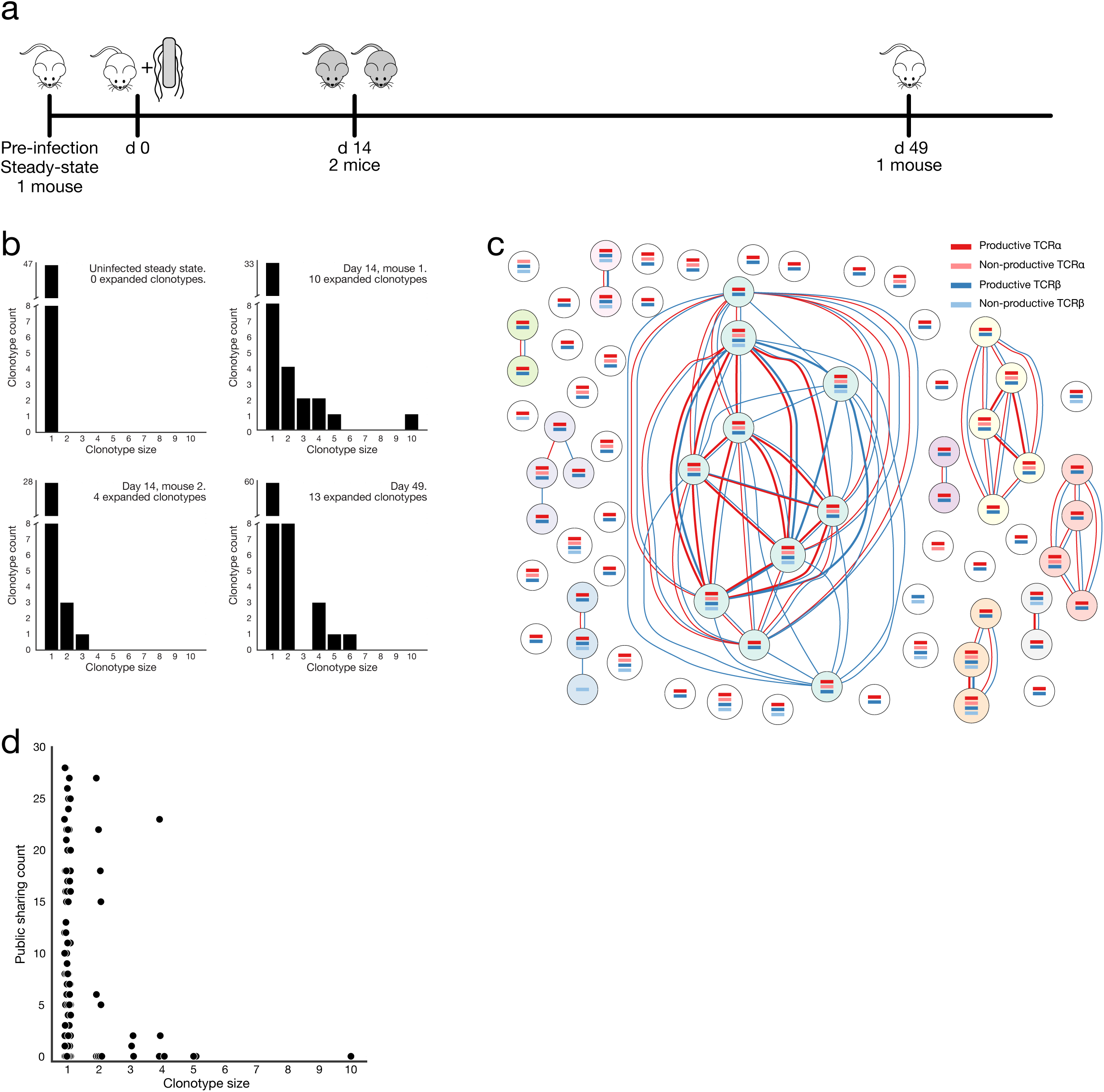
Assessment of clonal CD4^+^ T cell expansion during *Salmonella typhimurium* infection. (**a**) Schematic of timeline for *Salmonella* infection experiment. (**b**) Distribution of expanded clonotypes within splenic CD4^+^ T cell populations analysed by single-cell RNA-seq. The x-axis indicates the number of cells within the expanded clonotypes whilst the y-axis represents the number of clonotypes of each size. (**c**) Clonotype network graph from day 14, mouse 1. Each node in the graph represents an individual splenic CD4^+^ T lymphocyte. Coloured bars within the nodes indicate the presence of reconstructed TCR sequences that were detected for each cell. Dark coloured identifiers are productive, light coloured are non-productive. Red edges between the nodes indicate shared TCRα sequences whilst blue edges indicate shared TCRβ sequences. Edge thickness is proportional to the number of shared sequences. (**c**) Relationship between clonotype size and level of TCRβ CDR3 publicity. Reconstructed TCRβ nucleotide sequences from every cell analysed across all mice were translated and the CDR3 amino-acid sequences extracted. For each CDR3 sequence, its level of sharing between mice was determined from Madi *et al*^41^.

Taken together, these data indicate that our method accurately and sensitively determines the sequences of recombined and expressed TCR loci within individual T cells from single-cell RNA-seq data.

### Clonal expansion of CD4^+^ T lymphocytes during *Salmonella* infection

We demonstrated an application of our approach by investigating in detail the CD4^+^ T lymphocyte clonotypes present within the spleens of mice prior to, during or after a non-lethal infection with *Salmonella typhimurium. S. typhimurium* elicits a strong type-1 CD4^+^ T cell response and is widely used as murine model of enteric fever. Bacterial clearance and protective immunity require production of interferon-γ by Th1-type CD4^+^ T cells, and genetic defects in Th1-related signaling pathways have been shown to cause a predisposition to infection^33^,^34^. We analysed effector cells (CD44^high^CD62L^Low^) at day 14 when their relative abundance is close to its maximum, and memory cells (CD44^high^CD62L^Low^CD127^high^) at day 49 when the infection has been resolved^35^ (**Supplementary Fig. 1)**.

Analysis of the TCR sequences present within the splenic CD4^+^ T cells found 12 cells that expressed a productive TCRα recombinant comprising TRAV11 (Valpha14) and TRAJ18 (Jalpha18) segments (**Supplementary Table 1**). This gene combination is typical of invariant natural killer T (iNKT) cells^36^. Although there was variation between the exact junctional nucleotides within these TCRα sequences, each encoded exactly the same CDR3 amino acid sequence (CVVGDRGSALGRLHF), the first eight nucleotides of which were also previously reported as the most common junctional sequence from eight iNKT hybridomas^37^. Furthermore, 11 out of 12 cells expressing an iNKT-like TCRα chain also expressed a productive beta chain with a V gene from the limited repertoire observed previously in iNKT cells^36^.

The discovery of these iNKT cells within our analysed populations demonstrates one aspect of the utility of considering paired TCR sequences within individual cells under study. We excluded NK1.1^+^ cells during T cell purification but it is known that a CD4^+^NK1.1^-^ population of iNKT cells exists within the spleen^38^ and that NK1.1 is downregulated on NKT cells following *Salmonella* infection^39^. iNKT cells are best identified by their TCR specificity using CD1d tetramers loaded with α-GalCer^40^. Here, we show that we can identify such cells without requiring additional staining during cell-sorting procedures. Since we wished to focus on conventional MHC-restricted CD4^+^ T cells, we excluded the 12 iNKT cells from further analyses.

We compared recombinant identifiers between all cells to find cases where multiple cells expressed TCR genes with exactly the same nucleotide sequence. These cells are examples of clonal expansion whereby multiple members of the clonotype all derive from a single ancestor. We found no TCR sharing between cells from different mice nor between cells from within the uninfected mouse (**Fig. 3b**, **Supplementary Fig. 4**, **Supplementary Table 1**). The huge potential diversity of TCR nucleotide sequences means that it is extremely unlikely that a small number of CD4^+^ T cells sampled from a single steady-state mouse will exhibit evidence of TCR sharing. It is also equally unlikely that identical TCR nucleotide sequences will be found in cells from different mice.

**Figure 4.**
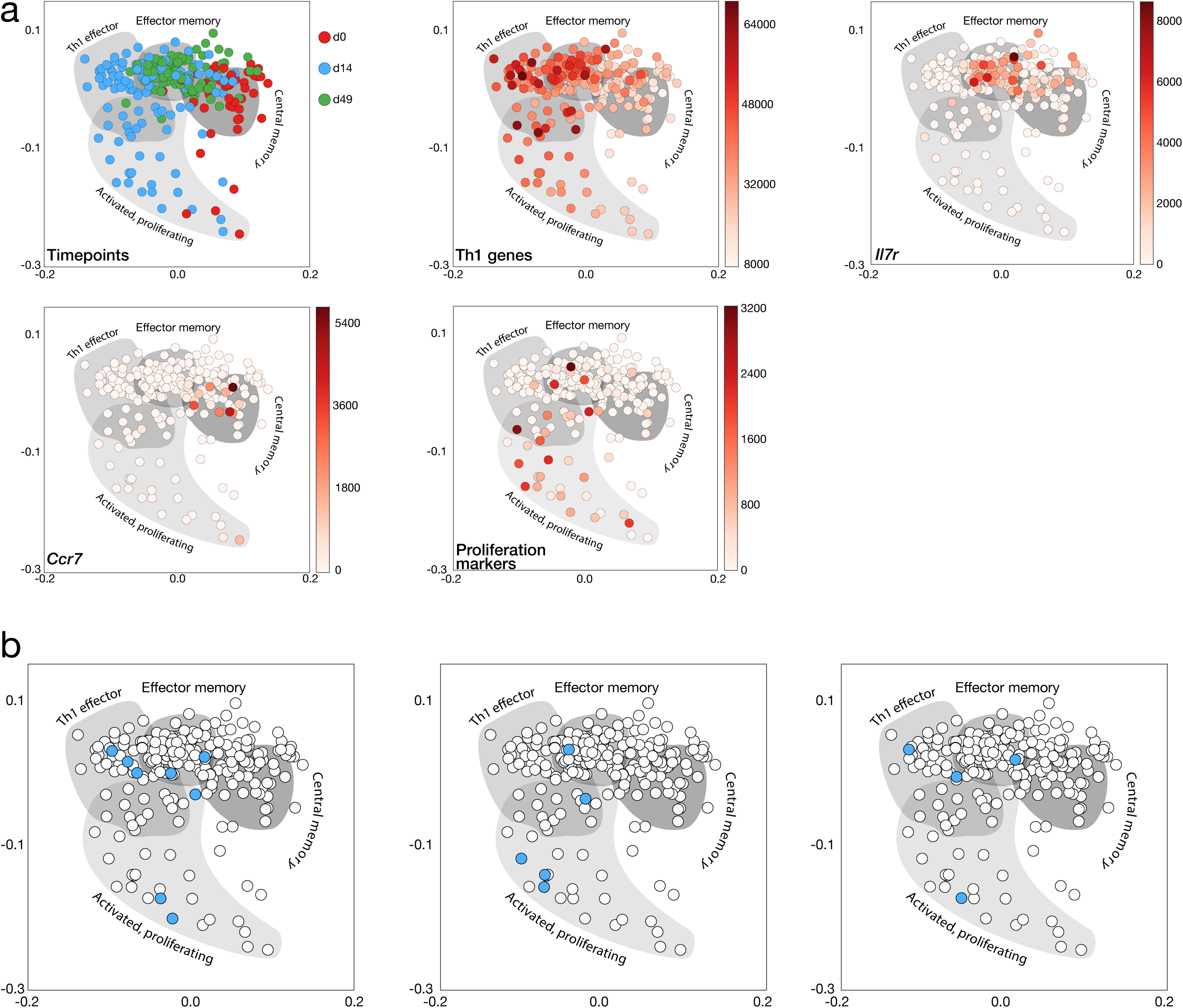
Distribution of expanded clonotypes throughout the Th1 response to *S. typhimurium* infection. (**a**) Dimensionality reduction of single-cell gene expression data by independent component analysis (ICA). Each single CD4^+^ T cell is plotted in reduced two-dimensional space according to its gene expression profile. Points are colored according to the timepoint from which they were sampled or according to their expression of marker genes indicative of their phenotype. Where the expression of a set of genes (Th1 genes and proliferation markers) is plotted, this is the sum of TPM values for the genes within the set. (**b**) Clonotype distribution in gene-expression space. Three representative expanded clonotypes from day 14 mouse 1 are shown as blue points on top of all other cells within the gene expression space.

We did see evidence of clonotype expansion within activated CD4^+^ T lymphocytes from each mouse at day 14 of *Salmonella* infection as well as within the effector memory cells from the mouse at week 7 post-infection (**Figs. 3b, 3c** and **Supplementary Figs. 5–7**. **Supplementary Table 1**). TCR sequences within expanded clonotypes from these mice are likely to be specific for *Salmonella* antigens and to be associated with clonal expansion of their T cells upon activation during the immune response to infection.

Importantly, we observe multiple cells that share all their detected recombinant sequences including those that are non-productive. This indicates that our method is detecting the correct combinations of TCR recombinants within the cells. We detected cells that share TCRβ sequences but have different TCRα sequences. This can be expected given that developing T lymphocytes in the thymus first perform recombination at the TCRβ locus and undergo proliferation prior to recombining their TCRα loci. Cells generated from a single progenitor by this proliferative expansion will all have the same TCRβ recombinant but will each randomly generate a different α recombinant prior to continuing their maturation and entering the periphery.

It should be noted that there is no evidence of contamination across microfluidics chip capture or harvest sites or adjacent wells in the 96-well plates used (**Supplementary Fig. 8**). Again, this implies that the observed sequence sharing represents a genuine biological phenomenon.

We compared the CDR3 amino acid sequences that we found within our data with those described in previous work that identified different degrees of CDR3 sharing between mice and classified sequences as ‘public’ or ‘private’^41^ (**Fig. 3c**). Public sequences were shared between the majority of mice whilst private sequences were only observed in very few mice. CDR3 sequences from expanded clones appear to be less likely to be public, a finding that agrees with the previous conclusion that pathogen-responsive CDR3s are more private.

### Distribution of clonally expanded cells between CD4^+^ T cell functional states

Single-cell RNA-seq allows cells to be classified according to their gene expression profiles. Our ability to accurately determine the paired TCR sequences within each single cell allows us to explore how cells that are all derived from a single ancestor in an expanded clonotype are distributed between transcriptional states. To demonstrate this, we quantified gene expression within each single CD4^+^ splenic T cell and performed independent component analysis (ICA) to reduce the gene expression space to two dimensions (**Fig. 4a**).

After filtering for genes expressed in at least three cells, we were able to use 14,889 informative genes for ICA. This is a great deal larger than the 17 phenotyping genes that were used in a previous PCR-based approach to determining clonality and cell fate^12^. We determined the phenotype of the cells within the reduced gene expression space by analysing the expression of 259 genes that indicate a Th1 cell fate^42^, *Il7r* (CD127) which is indicative of effector-memory T cells^43^, *Ccr7* (a marker of central-memory T cells)^44^ and a set of seven genes that are expressed in proliferating cells^45^ (**Fig 4a**).

Expression of these genes and modules of genes allowed us to separate the cells into four populations: activated proliferating cells that are differentiating to the Th1 fate, mature differentiated Th1 effector cells, effector memory-like cells and central memory-like cells. Cells from the steady-state mouse are mostly central memory-like, cells from the mouse at day 14 have an activated or Th1 effector phenotype whilst cells from day 49 (sorted to be CD127^high^, a marker of effector memory fate) are found in the effector-memory region of the ICA gene expression space.

We then determined the distributions of expanded clonotypes within the reduced gene-expression space (**Fig 4b** and **Supplementary Figs. 9-11**). Cells derived from the same progenitor can be seen throughout the activated differentiating, Th1 effector and effector-memory populations. This suggests that, after activation by binding to a *Salmonella* antigen–MHC complex, the progeny of a particular CD4^+^ T cell differentiate to the effector and memory subtypes at varying rates leading to the asynchronous populations that we observe here. In other words, the members of one clonotype can be spread across the full spectrum of proliferation and differentiation states that occur during the *Salmonella* response.

## DISCUSSION

Here, we present a method for the determination of paired T cell receptor sequences from individual T lymphocytes achieved solely by analysis of standard single-cell RNA-seq datasets without the need for alterations to the RNA-seq protocol. Our method is as sensitive as the best-performing PCR-based method^12^ for determining paired, productive α and β chains. For the detection of cases where both alleles of the β locus have been recombined, our method achieves better sensitivity than the PCR-based method. This is likely due to RNA-seq not being subject to PCR biases so that multiple recombinants within a cell are detected independently. Furthermore, our method is easy to adapt to any species for which annotated TCR gene sequences are available without the need for design and lengthy optimization of large numbers of multiplexed PCR reactions. We also fully expect our method to be easily adapted to the study of the analogous B cell receptor/antibody sequences within B lymphocytes.

Reconstruction of TCR sequences from single-cell RNA-seq datasets means that the information about cell lineage and antigen specificity is obtained alongside the comprehensive transcriptomic identity of the cells. This provides us with the opportunity to assess the cells’ phenotypes and to perform clustering analysis using orders of magnitude more genes than existing PCR-based approaches. RNA-seq also obviates the need for *a priori* knowledge of phenotyping genes of interest. This will permit the discovery of novel or poorly-characterised phenotypic subtypes in conjunction with the analysis of their TCR sequences. Our method will work with any scRNA-seq protocol that produces reads from full-length cDNA. This will become increasingly valuable as higher-throughput scRNA-seq methods are developed and applied to T and B lymphocytes.

Having assessed the performance of our method and validated it using an alternative technique, we demonstrated its ability to analyse the distribution of murine T cell clones between phenotypically different CD4^+^ T cell populations during *S. typhimurium* infection. Challenge with *Salmonella* is known to cause large clonal expansion of responsive T cells within the spleen^35^,^46^,^47^ and we were able to observe expanded clonotypes within our samples. Full transcriptomic quantification of gene expression allowed us to perform fine-grained determination of the various cell states within the Th1 response. We showed that members of a single expanded clonotype can be found at different stages of effector T cell activation and also within the population of memory cells. This demonstrates that T cells derived from a single progenitor can exhibit divergent fates at the same time within a single mouse.

To enable the widespread use of our method by researchers who perform single-cell RNA-seq on lymphocyte populations we have made the TraCeR tool and associated documentation freely available for download. A combined knowledge of T cell clonal dynamics, TCR specificity and detailed transcriptional phenotype is likely to be of great use in the study of T cell responses to infection, auto-antigens or vaccination and will provide insights into both pathogenic mechanisms and therapeutic approaches.

## METHODS

### Ethics statement

Mice were maintained under specific pathogen-free conditions at the Wellcome Genome Campus Research Support Facility (Cambridge, UK). These animal facilities are approved by and registered with the UK Home Office. Animals were sacrificed by approved animal technicians in accordance with Schedule 1 of the Animals (Scientific Procedures) Act 1986. Oversight of the arrangements for Schedule 1 killing was performed by the Animal Welfare and Ethical Review Body of the Wellcome Genome Campus.

### Cell preparation

Female C57BL6/N mice aged 6-8 weeks were infected intravenously with 0.2 ml Salmonella Typhimurium M525 containing 5 × 10^5^ CFU of bacteria in sterile phosphate buffered saline (PBS, Sigma-Aldrich). At day 14 or 49 post infection (p.i.) mice were sacrificed with spleens and livers being harvested. Bacteria were enumerated from the livers by serial dilution and plating onto agar plates (Oxoid) to confirm levels of infection. Single-cell suspensions were prepared by homogenising spleens through 70 μm strainers and lysing erythrocytes. Following incubation with CD16/CD32 blocking antibody, the cells from day 14 p.i. were stained with titrated amounts of fluorochrome conjugated antibodies for CD44(FITC), CD25(PE), CD62L(PE-CF594), TCRβ(PerCP-Cy5.5), CD69(APC), CD8α(APC-H7), CD161(BV421), and CD4(BV510). The cells from day 49 p.i. were stained with antibodies for CD44(FITC), CD127(PE), CD62L(PE-CF594), TCRβ(PerCP-Cy5.5), CD161(APC), CD8α(APC-H7), CD4 (BV510), and Sytox Blue viability stain. Cell sorting was performed using a BD FACSAria II instrument using the 100 micron nozzle at 20 psi using the single cell sort precision mode. The cytometer was set up using Cytometer Setup and Tracking beads and compensation was calculated using compensation beads (for antibodies, eBioscience UltraComp) and cells (for Sytox Blue) using automated software (FACSDiva v6).

### Single-cell RNA-sequencing and gene expression quantification

Capture and processing of single CD4^+^ T cells was performed using the Fluidigm C1 autoprep system. Cells were loaded at a concentration of 1700 cells μl^−1^ onto C1 capture chips for 5-10 μm cells. ERCC (External RNA Controls Consortium) spike-in RNAs (Ambion, Life Technologies) were added to the lysis mix. Reverse transcription and cDNA preamplification were performed using the SMARTer Ultra Low RNA kit (Clontech). Sequencing libraries were prepared using Nextera XT DNA Sample Preparation kit with 96 indices (Illumina), according to the protocol supplied by Fluidigm. Libraries were pooled and sequenced on Illumina HiSeq2500 using paired-end 100 base reads.

Reads were mapped to the *Mus musculus* genome (Ensembl version 38.70) concatenated with the ERCC sequences, using GSNAP^48^ with default parameters. Gene-specific read counts were calculated using HTSeq^49^. Sixty-eight cells (out of 352) with detected transcripts for fewer than 2000 genes, or with more than 10% of measured exonic reads corresponding to genes coded by the mitochondrial genome, were excluded from further analyses.

### Reconstruction and analysis of TCR sequences from RNA-seq data

Synthetic genome files were separately created for the TCRα and TCRβ chains. To generate these fasta files, nucleotide sequences for all mouse V and J genes were downloaded from The International ImMunoGeneTics information system^50^ (IMGT, www.imgt.org). Every possible combination of V and J genes was generated for each TCR locus such that each combination was a separate sequence entry in the appropriate synthetic genome file. Ambiguous N nucleotides were introduced into the junction between V and J genes in each sequence entry to improve alignments of reads that spanned diverse junctional sequences. Seven N nucleotides were used in TCRβ combinations whilst one N nucleotide was used in the TCRα combinations. V gene leader sequences are not well annotated within IMGT and so 20 N nucleotides were added at the 5′ end of the V sequence to permit alignment of sequencing reads that included the leader sequence.

TCRα or TCRβ constant region cDNA sequences were downloaded from ENSEMBL and appended to the 3′ end of each combined sequence to permit alignment of reads that ran into the constant region. The full-length TCRα constant region was used whilst the only the first 259 nucleotides of the TCRβ constant gene were used since these are identical between both *Trbc* homologs that are found within the mouse genome. The synthetic genomes used in this work can be found alongside the other tools at www.github.com/teichlab/tracer.

RNA-seq reads from each cell were aligned against each synthetic genome independently using the Bowtie 2 aligner^51^. Bowtie 2 is ideal for alignment against the synthetic genomes because it can align against ambiguous N nucleotides within a reference and also introduce gaps into both the reference and read sequences. This allows it to align reads against the variable junctional regions. We used the following Bowtie 2 parameters with low penalties for introducing gaps into either the read or the reference sequence or for aligning against N nucleotides ‘--no-unal -k 1 --np 0 --rdg 1,1 --rfg 1,1’. For each chain, separately, we used the reads aligning to the synthetic genome as input to the Trinity RNA-seq assembly software^27^ using its default parameters.

V, D and J gene sequences downloaded from IMGT were used to generate appropriate databases for use with IgBLAST^28^. Contigs assembled by Trinity were used as input to IgBlast and the resulting output text files were processed with a custom parsing script. Contigs were classed as representing TCR sequences if they contained gene segments from the correct locus (ie TCRα genes for TCRα contigs) and if their reported V and J alignments had E-values below 5 × 10^−3^. If multiple contigs within the same cell represented the same recombinant sequence, these were collapsed so that the sequence was only represented once in the cell for subsequent analyses. In some cases where two contigs derived from the same original sequence but one was shorter than the other, IgBLAST assigned different V sequences if the shorter sequence did not provide sufficient information to distinguish between highly similar genes. This typically occurred with V genes that were part of the evolutionary expansion events that caused gene duplication and triplication within the TCRα locus^29^. In these cases, the sequences were collapsed into a single assignment that used the results from the longest contig. The IgBLAST results for the TCR sequences within each cell were then reduced to an identifying string consisting of the V gene name, the junctional nucleotide sequence and the J gene name (eg. TRBV31_AGTCTTGACACAAGA_TRBJ2-5) which was used for comparisons between sequences within other cells.

It is important to determine whether a particular TCR mRNA sequence is productive and therefore able to be translated to produce a full-length TCR polypeptide chain. To do this for the reconstructed TCR sequences we first converted them to entirely full-length sequences by using full-length V and J gene sequences from IMGT appropriate to the gene segments assigned by IgBLAST. Since TCRs do not undergo somatic hypermutation we can make the assumption that variations between the RNA-seq derived sequences and the reference sequences outside the junctional region are due to PCR and/or sequencing errors and so can be ignored. We check that these full-length sequences are in the correct reading frame from the start of the V gene to the start of the constant gene and that they lack stop codons. If this is the case, the sequence is classed as productive. For analysis of CDR3 amino-acid sequences we translate the productive recombinants and define the CDR3 as the region flanked by the final cysteine residue of the V gene and the conserved FGXG motif in the J gene as previously described^23^.

Expression levels of the TCR genes found within a cell were quantified by appending that cell’s full-length recombinant sequences to a file containing the entire mouse transcriptome (downloaded from http://bio.math.berkeley.edu/kallisto/transcriptomes/) and then using this file for the generation of an index suitable for use with the pseudoalignment-based Kallisto algorithm^31^. This index was then used with the RNA-seq reads for the cell as input for Kallisto in quantification mode to calculate transcripts per million (TPM) values for each TCR sequence. If a cell was assigned more than two recombinant sequences for a particular locus, the sequences were ranked by their TPM values and the two most highly-expressed were used for further analyses. Kallisto’s speed in constructing indices and performing expression quantification makes it ideal for this task.

After assignment of TCR sequences to each cell within an experiment, we used custom Python scripts to compare the recombinant identifiers present in each cell to find cases where multiple cells contained the same identifier. These analyses were used to generate network graphs where each node in the graph represents a single cell and edges between the nodes represent shared TCR sequences.

The analyses described above are performed by our tool, TraCeR which is freely available at www.github.com/teichlab/tracer.

### PCR-based sequencing of TCR sequences

Primers were designed to amplify all possible recombined TCR sequences from both the TCRα and TCRβ loci (all sequences can be found in **Supplementary Table 3**). Two constant region primers were designed to be complementary to the *Trac* or *Trbc* genes close to their 5′ ends. Sets of primers complementary to all TCRα and β V gene sequences downloaded from IMGT were also designed. Primers were designed to regions of homology between V genes and included degeneracy where appropriate so as to minimise the number of primers required. In total, 34 TCRα and 31 TCRβ primers were used. All primers were designed with a T_m_ of 71–73 °C. All V gene primers were designed with the sequence ACACTCTTTCCCTACACGACGCTCTTCCGATCT at their 5′ end to allow amplification by the Illumina PE 1.0 primer (AATGATACGGCGACCACCGAGATCTACACTCTTTCCCTACACGACGCTCTTCCG ATCT) while the constant region primers were designed with the sequence TCGGCATTCCTGCTGAACCGCTCTTCCGATCT at their 5′ end so that they could be amplified by barcoding primers containing a unique 11 nt index sequence (**Supplementary Table 3**). The barcoding primers also contain the Illumina PE 2.0 sequence.

Full-length (oligo-dT primed) cDNA produced from single-cells by the C1 system (Fluidigm, USA) was used as template in two PCR reactions, one for each TCR locus. 0.4 µl of cDNA were used in each reaction along with each V primer at 0.06 µM and the constant primer at 0.3 µM. Phusion DNA polymerase (NEB, USA) was used to perform the amplification in 25 µl final volume. The cycling conditions for this step were 98 °C 30 s; 98 °C 10 s, 60 °C 10s, 72 °C 30 s × 16 cycles; 72 °C 5 min. 4 °C. A 1 µl aliquot of the first reaction was used as template in a second PCR amplification, again using Phusion in a 25 µl reaction volume. Here, the Illumina PE 1.0 primer was used with a barcoding primer unique for each cell and each primer was at 0.4 µM. The cycling conditions for this step were 98 °C 30 s; 98 °C 10 s, 58°C 10s, 72 °C 30 s × 16 cycles; 72 °C 5 min. 4 °C. PCR products of the correct size for sequencing were purified using 0.7 volumes of AMPure XP beads (Beckman Coulter) according to the manufacturer’s instructions. Purified products were pooled and submitted to the Wellcome Trust Sanger Institute (WTSI) Sequencing Facility for sequencing using a MiSeq (Illumina) with 250bp paired-end reads.

### Processing PCR data

Reads generated by MiSeq sequencing of PCR products were de-multiplexed by the WTSI Sequencing Facility according to their barcode sequences. Reads were then trimmed to remove low-quality regions and adapter sequences using TrimGalore (http://www.bioinformatics.babraham.ac.uk/projects/trim_galore/). The TCR-targeted PCR primers were designed to provide amplicons short enough such that the forward and reverse paired reads would overlap upon sequencing enabling read pairs to be merged using FLASH^52^. Merged read sequences were then filtered to remove those under 200 nucleotides in length to remove artefactual sequences. Following this step, read sequences for each cell were subsampled where necessary such that there were 50,000 sequences or fewer from each cell. This reduced the computational time and requirements for the next stage whilst still providing sufficient information about the sequences present. As described previously^12^, we assumed that sequences from an individual cell that had at least 95% sequence identity were derived from the same original cDNA sequence and so these were combined to generate a consensus sequence. The consensus sequences for each cell were analysed by IgBLAST to find sequences that represented recombined TCRs and the number of sequencing reads supporting each TCR were used to filter out background sequences that had few reads.

### Comparing PCR and RNA-seq data

For each cell, sequences derived from PCR analysis or reconstructed from RNA-seq data were trimmed to only include the regions assigned by IgBlast as containing V, D or J sequences. This removed any leader sequences or constant regions. Trimmed reconstructed RNA-seq sequences were aligned against the trimmed PCR-derived sequences in a set of pairwise comparisons using BLAST. If an alignment was reported, the number of mismatches across the entire alignment were counted, as were the number of mismatches between the nucleotides that encoded the CDR3 region (defined here as the 30nt following the end of the framework 3 region as annotated by IgBlast). If the CDR3 regions contained any mismatches, the alignment was classed as discordant, otherwise the two sequences were classed as concordant. Sequences from one method (RNA-seq or PCR) that did not align successfully with any sequence from the other method were classed as discordant.

### Gene expression quantification and dimensionality reduction

Genes were filtered to remove those expressed (TPM>1) in fewer than three cells. Dimensionality reduction of the remaining gene expression data was performed by independent component analysis (ICA) using the *FastICA* Python package.

For plotting gene expression for each cell within ICA space, 259 genes indicating a Th1-like fate and seven indicators of proliferation (*Mki67*, *Mybl2*, *Bub1, Plk1*, *Ccne1*, *Ccnd1*, *Ccnb1*) were taken from previous work^42^,^45^ and their expression levels (in TPM) were summed for each cell.

### Clonotype distribution within gene expression space

Cells that did not appear to be derived from the same progenitor (same TCRβ but differing TCRα chains) were removed from the expanded clonotype groups. Cells belonging to a particular expanded clonotype were then plotted within the ICA reduced gene expression space.

## ACKNOWLEDGEMENTS

We thank Valentine Svensson, Tzachi Hagai, Johan Henriksson and other members of the Teichmann laboratory for helpful discussions. We thank the Wellcome Trust Sanger Institute Sequencing Facility for performing Illumina sequencing and the Wellcome Trust Sanger Institute Research Support Facility for care of the mice used in these studies. This work was supported by European Research Council grant ThSWITCH (number 260507) and the Lister Institute for Preventative Medicine.

### AUTHOR CONTRIBUTIONS

MJTS conceived the project, designed the computational method, wrote the software, designed PCR sequencing primers, analysed data, generated figures and wrote the manuscript. TL and SC designed and performed the Salmonella experiments. TL performed cell collection and purification, generated scRNA-seq libraries, performed gene expression analyses, analysed data, generated figures and wrote the manuscript. VP performed PCR-based TCR sequencing experiments. AOS designed the cell sorting strategy, performed the sorting and generated figures. SAT and GD supervised work and wrote the manuscript.

## SUPPLEMENTARY TABLE TITLES

Supplementary Table 1 TCR sequences reconstructed from single-cell RNA-sequencing data.

Supplementary Table 2 Comparison between RNA-seq reconstruction and PCR-based detection of TCR sequences.

Supplementary Table 3 PCR primers used for TCR sequencing.

---

**Supplementary Figure 1.**
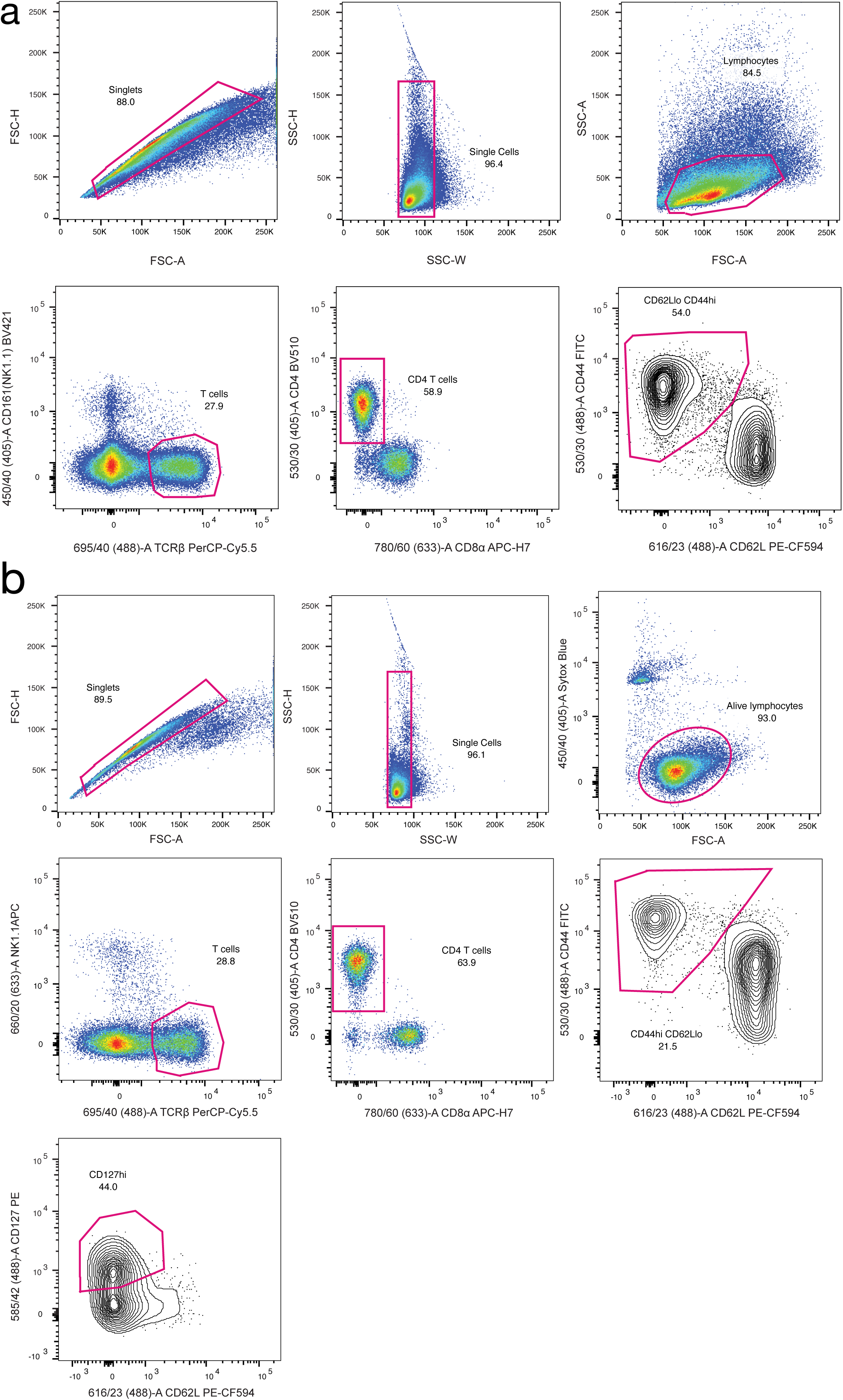
FACS strategy for (a) steady-state control and day 14 cells and (b) day 49 cells. In both cases, cells were sorted to be CD4^+^TCRB^+^NK1.1^-^CD44^+^CD62L^Low^. Additionally, cells from the mouse at day 49 were sorted to be CD127^High^.

**Supplementary Figure 2.**
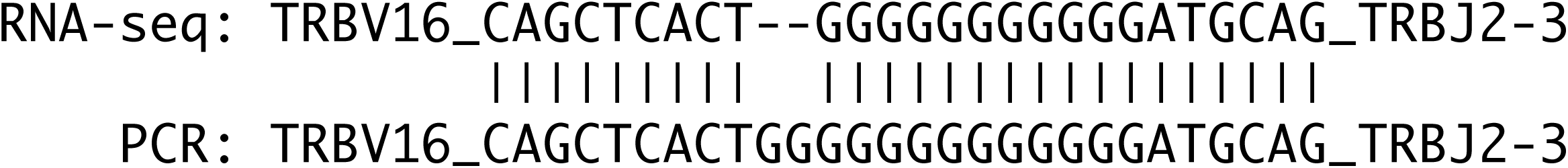
Discordancy between PCR and RNA-seq TCR sequence due to sequencing error. The TCR identifiers above were found in the same cell by RNA-seq and PCR. They differ solely by two G residues within the long homopolymeric G tract within the junctional region.

**Supplementary Figure 3.**
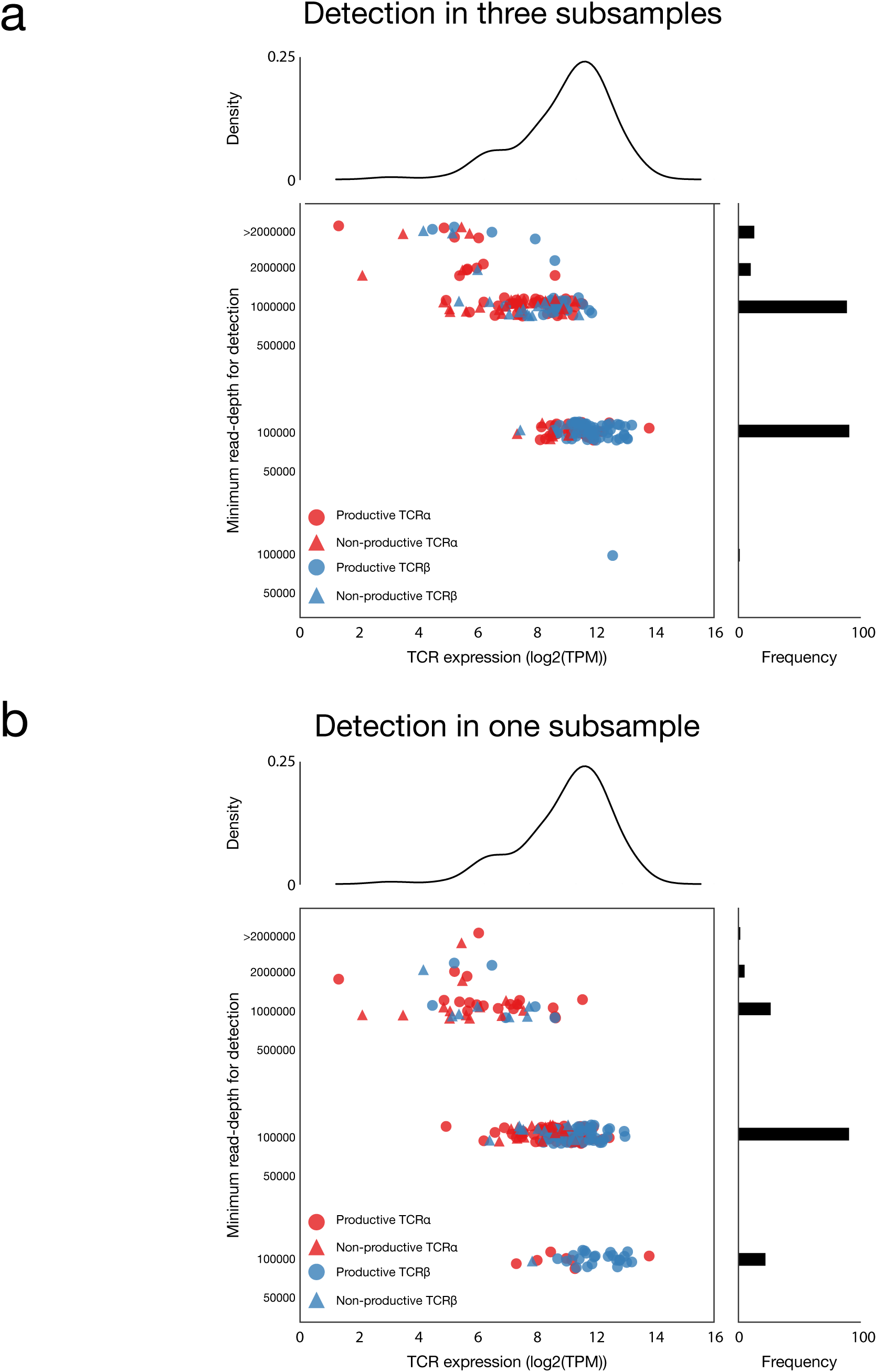
Sensitivity analysis of RNA-seq reconstruction. All single-cell datasets from day 14 mouse 1 were randomly subsampled three independent times to contain decreasing total read numbers followed by TCR reconstruction. Points representing each TCR sequence found in the full datasets are plotted according to their expression levels and the minimum total read depth required for detection in at least (a) three or (b) one out of three subsamples. For clarity, points are jittered about the y-axis.

**Supplementary Figure 4.**
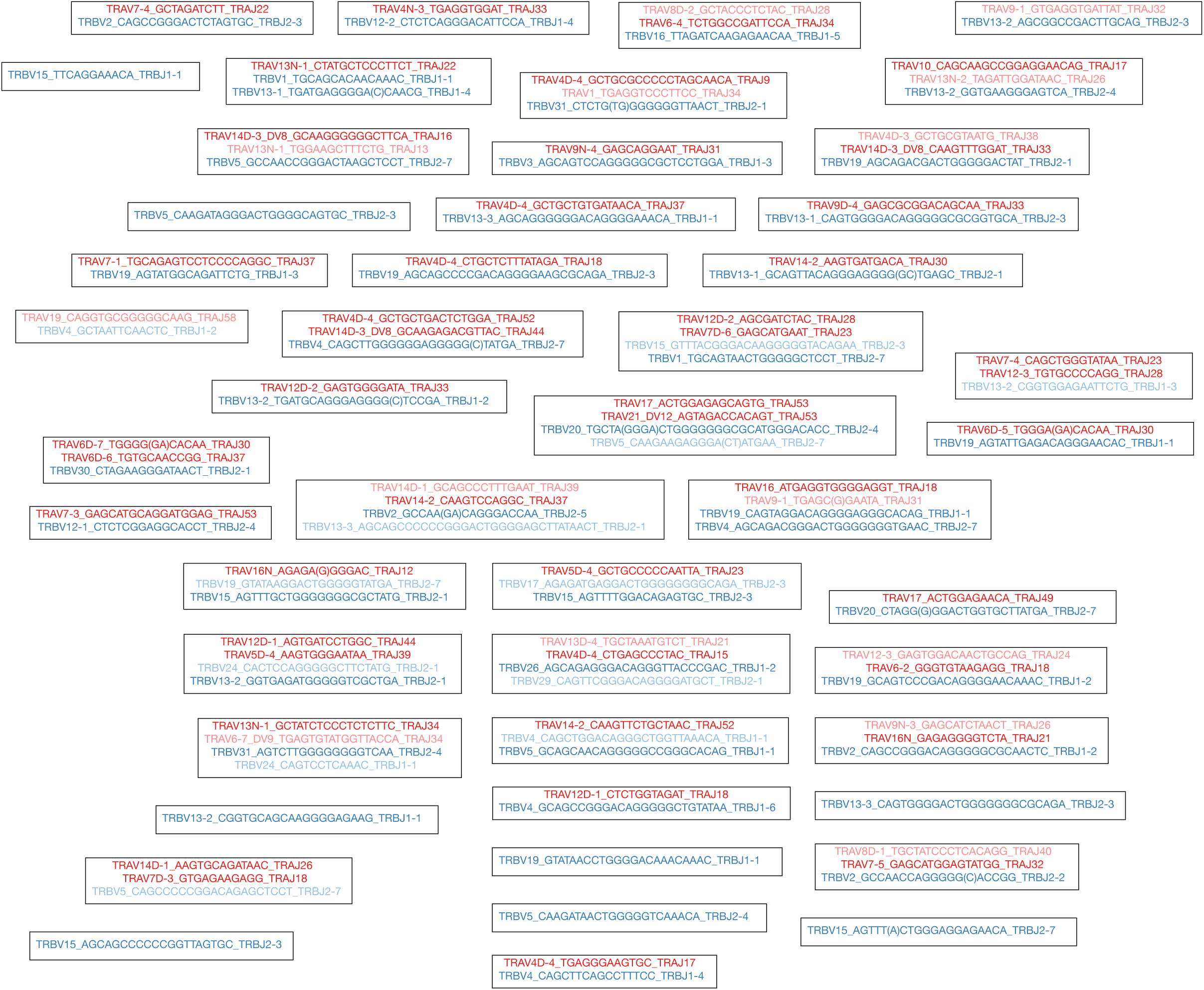
Clonotype network graph from steady-state uninfected mouse. Each node in the graph represents an individual splenic CD4+ T lymphocyte. Identifiers within the nodes indicate the reconstructed TCR sequences that were detected for each cell. Dark coloured identifiers are productive, light coloured are non-productive. The lack of edges between nodes in this graph indicates that no nodes share TCR sequences.

**Supplementary Figure 5.**
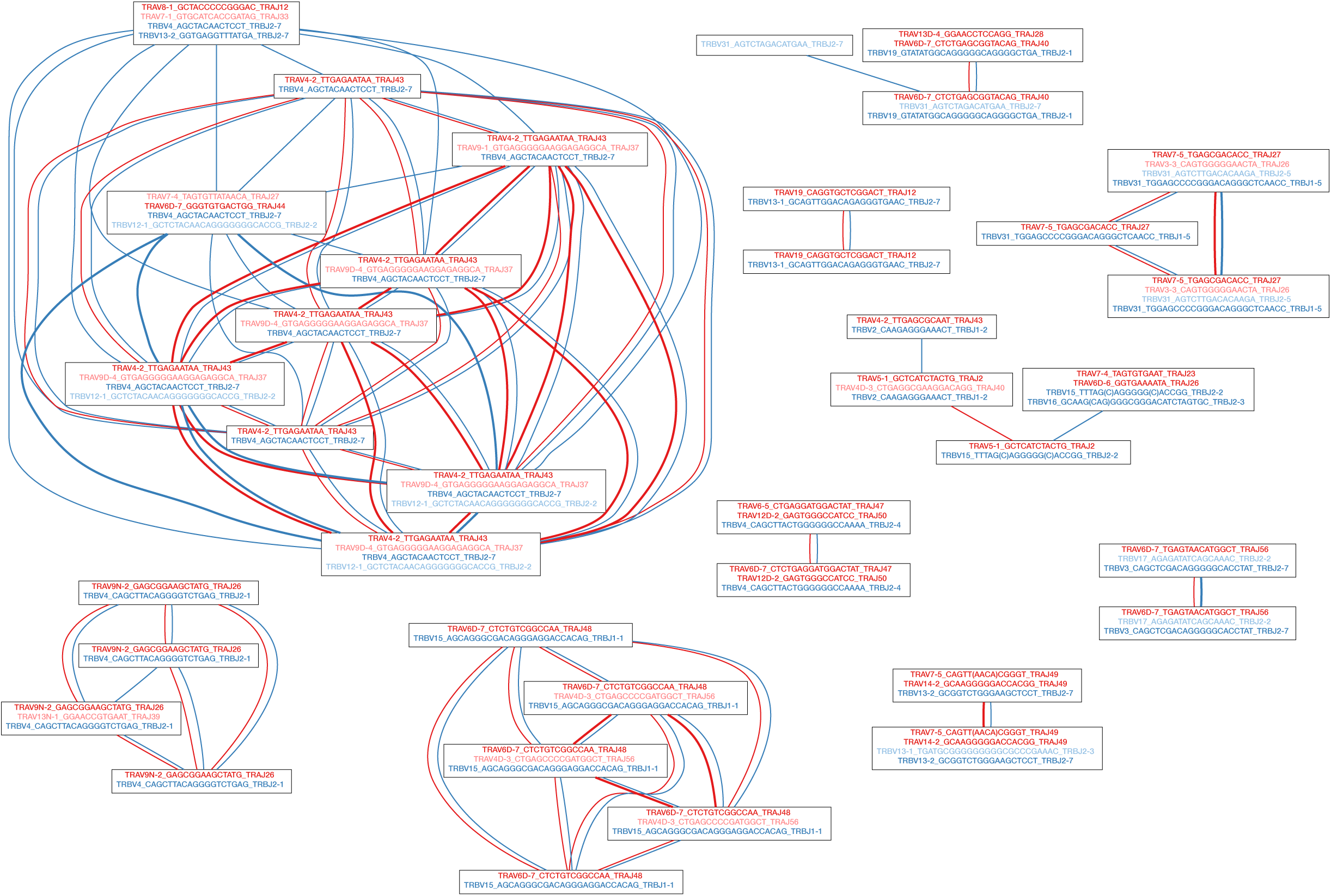
Clonotype network graph from day 14, mouse 1. Each node in the graph represents an individual splenic CD4+ T lymphocyte. Identifiers within the nodes indicate the reconstructed TCR sequences that were detected for each cell. Dark coloured identifiers are productive, light coloured are non-productive. Red edges between the nodes indicate shared TCRα sequences whilst blue edges indicate shared TCRβ sequences. Edge thickness is proportional to the number of shared sequences. For clarity, the 33 nodes without edges are not displayed.

**Supplementary Figure 6.**
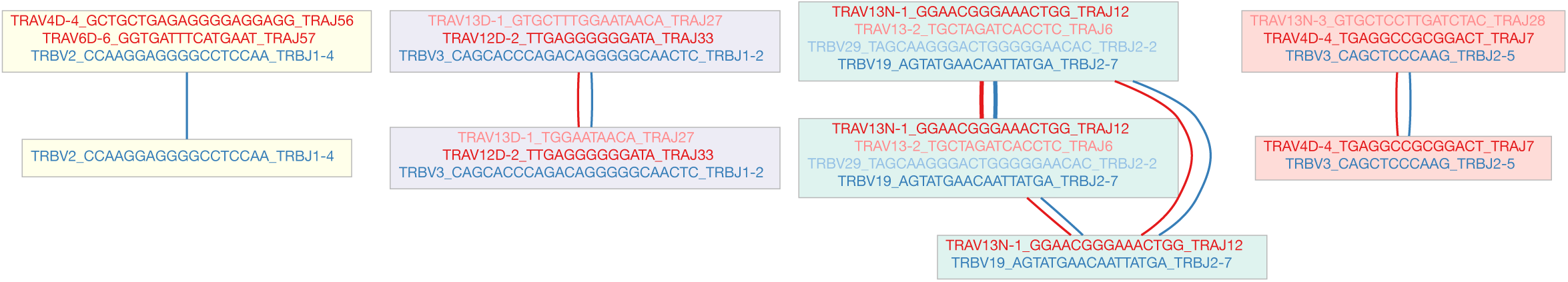
Clonotype network graph from day 14, mouse 2. Each node in the graph represents an individual splenic CD4+ T lymphocyte. Identifiers within the nodes indicate the reconstructed TCR sequences that were detected for each cell. Dark coloured identifiers are productive, light coloured are non-productive. Red edges between the nodes indicate shared TCRα sequences whilst blue edges indicate shared TCRβ sequences. Edge thickness is proportional to the number of shared sequences. For clarity, the 28 nodes without edges are not displayed.

**Supplementary Figure 7.**
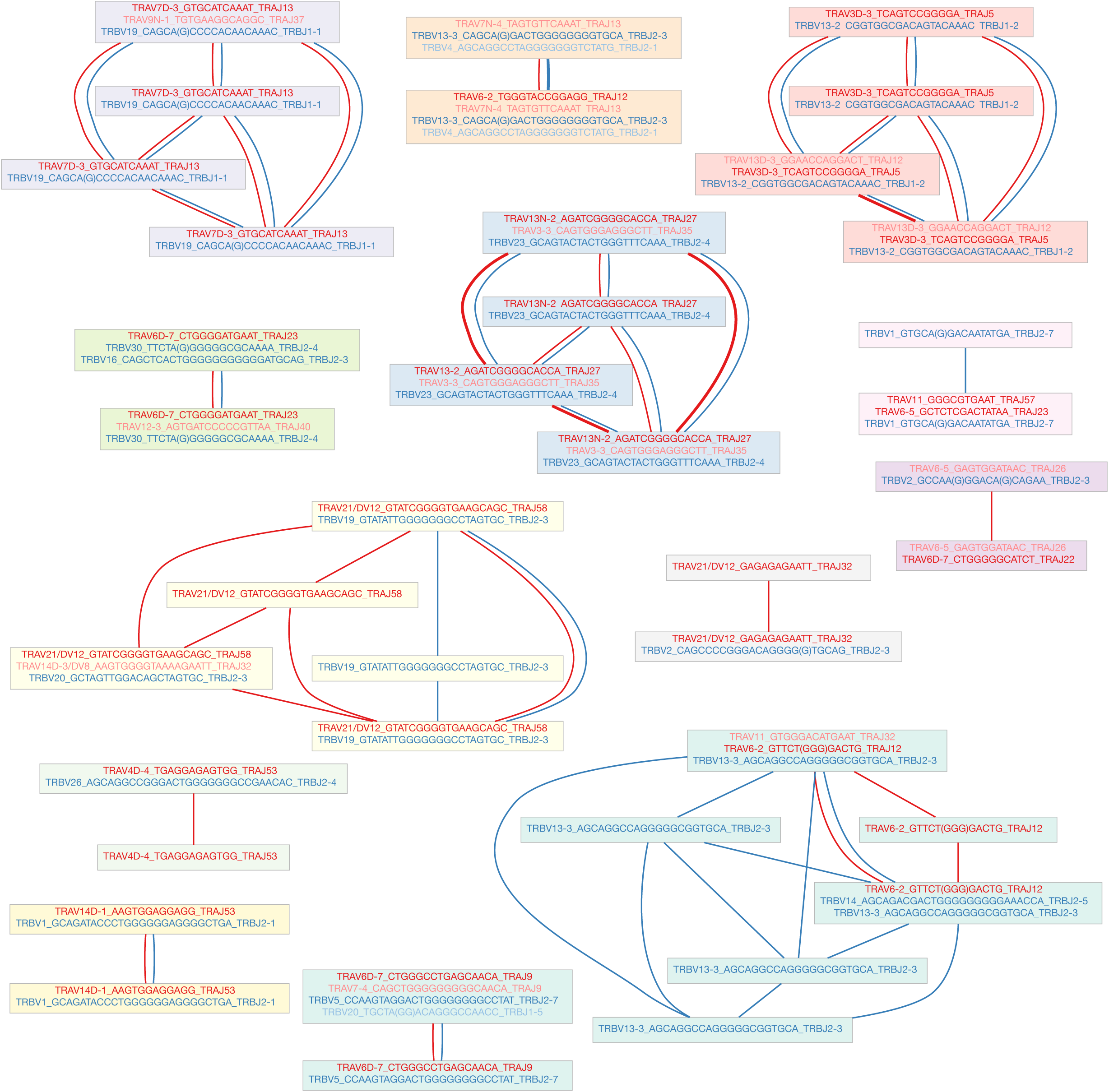
Clonotype network graph from day 49 mouse. Each node in the graph represents an individual splenic CD4+ T lymphocyte. Identifiers within the nodes indicate the reconstructed TCR sequences that were detected for each cell. Dark coloured identifiers are productive, light coloured are non-productive. Red edges between the nodes indicate shared TCRα sequences whilst blue edges indicate shared TCRβ sequences. Edge thickness is proportional to the number of shared sequences. For clarity, the 60 nodes without edges are not displayed.

**Supplementary Figure 8.**
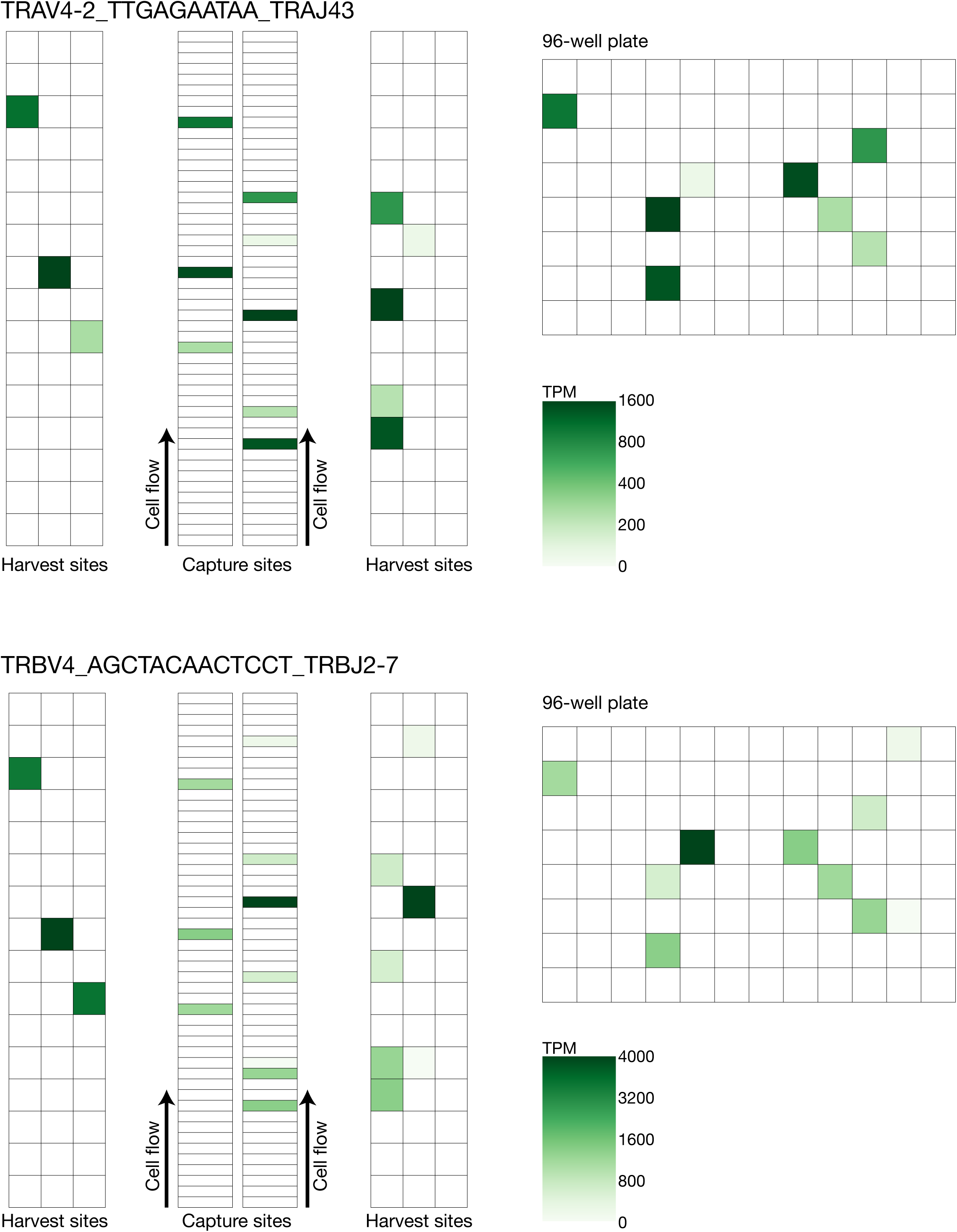
Distribution of shared TCR sequences within the Fluidigm C1 integrated fluidics circuit (IFC) and 96-well plate. TPM expression values of the two most highly-shared TCR sequences are shown within the C1 IFC capture sites, harvest sites and the resulting 96-well plate that contained the associated single cells.

**Supplementary Figure 9.**
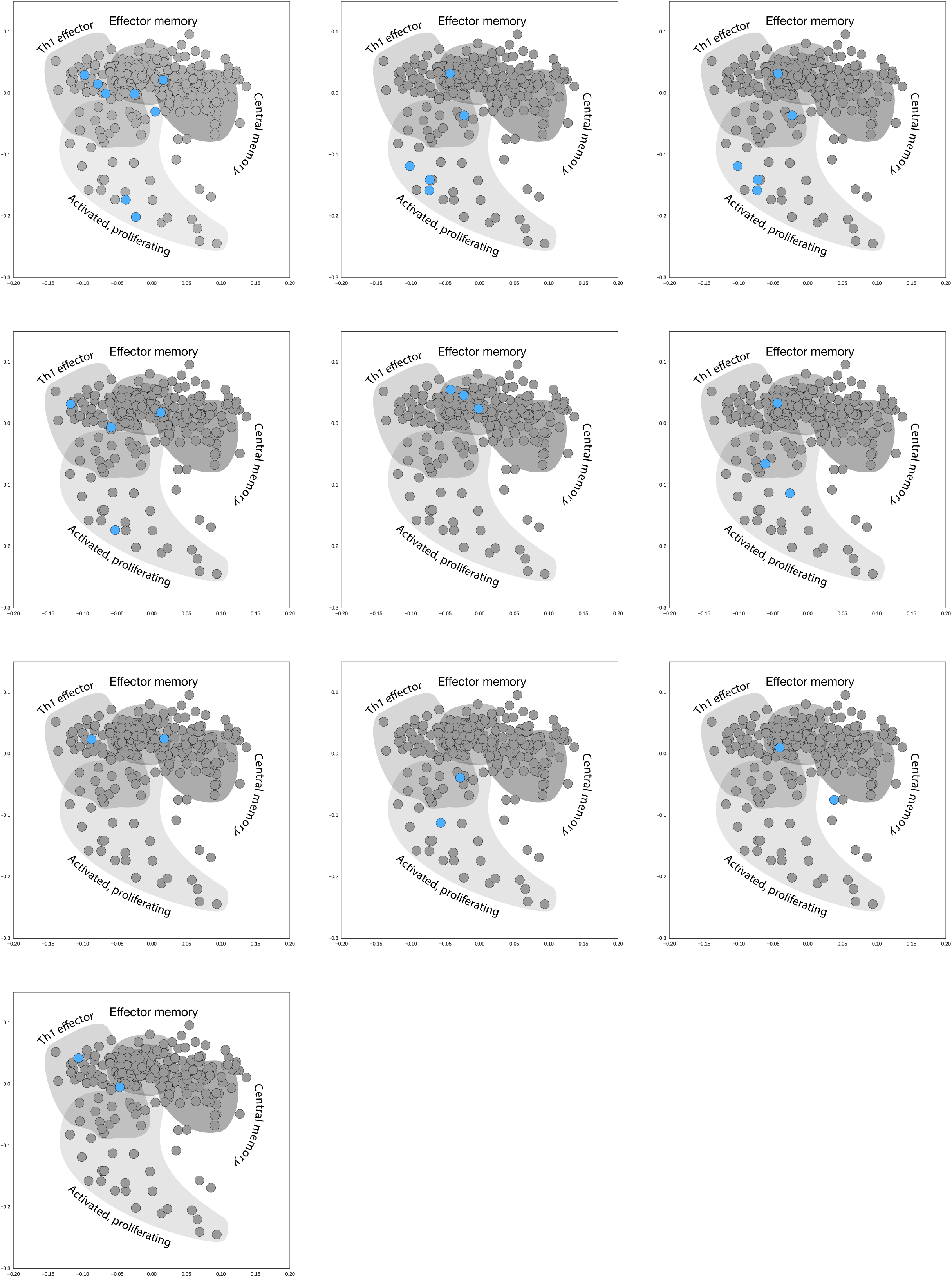
Clonotype distribution in gene-expression space. All clonotypes from day 14 mouse 1 are shown as blue points on top of all other cells within the gene expression space.

**Supplementary Figure 10.**
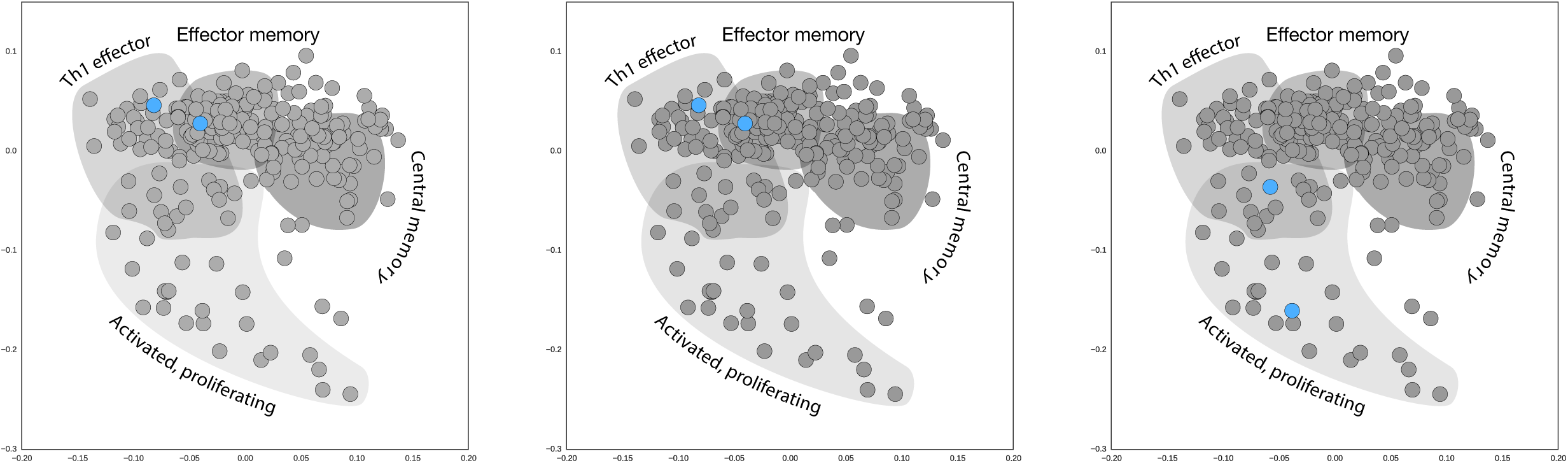
Clonotype distribution in gene-expression space. All clonotypes from day 14 mouse 2 are shown as blue points on top of all other cells within the gene expression space.

**Supplementary Figure 11.**
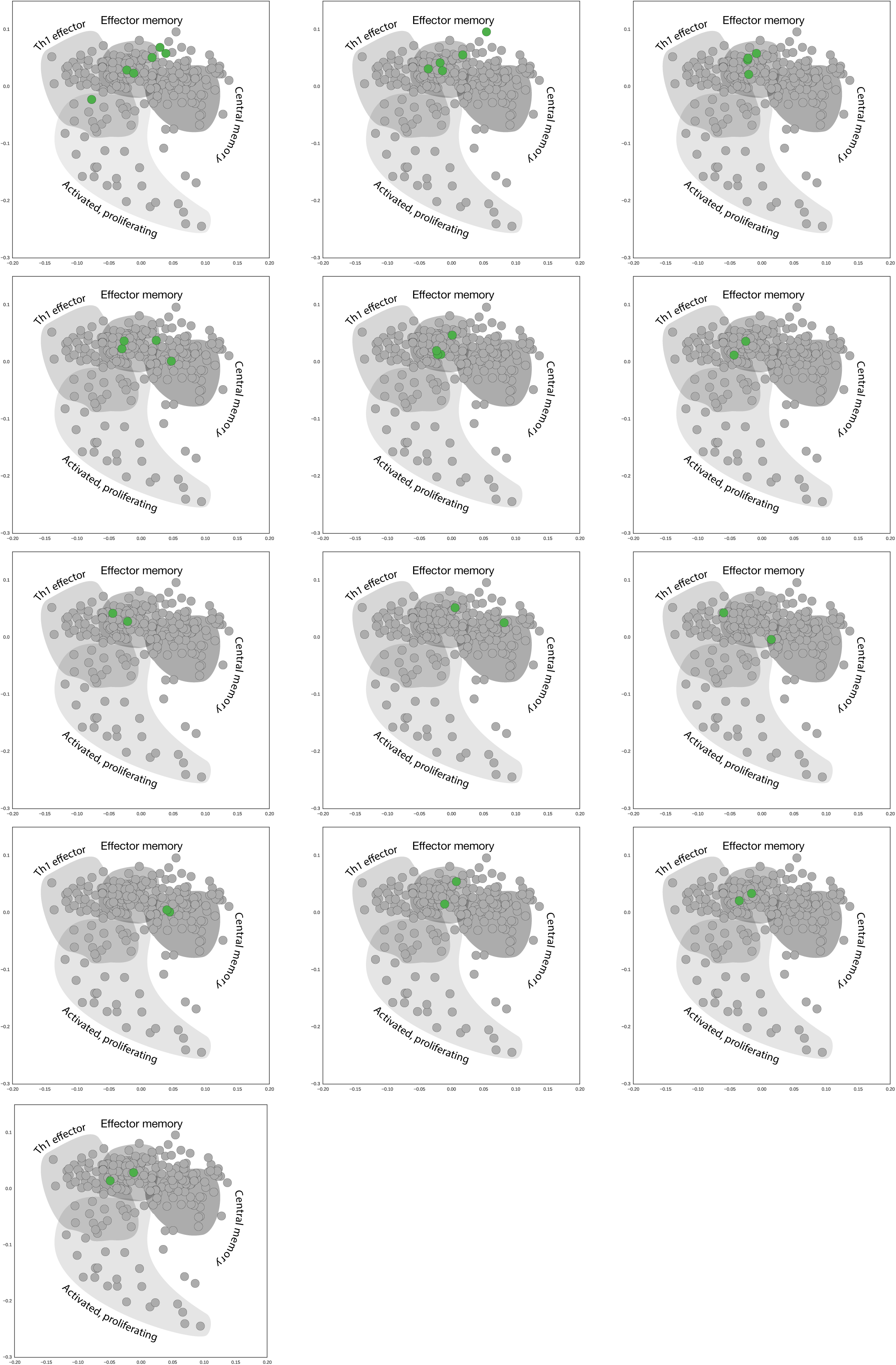
Clonotype distribution in gene-expression space. All clonotypes from the day 49 mouse are shown as green points on top of all other cells within the gene expression space.

